# Demographic and Historical Processes Influencing *Cochliomyia hominivorax* (Diptera: Calliphoridae) Population Structure across South America

**DOI:** 10.1101/2024.09.25.615064

**Authors:** Kelly da Silva e Souza, Letícia Chiara Baldassio de Paula, Ana Maria Lima de Azeredo-Espin, Tatiana Teixeira Torres

## Abstract

**Background:** This study investigates the genetic variability and population structure of *Cochliomyia hominivorax*, the New World screwworm fly. This study tested the hypothesis that the species exhibits a center-periphery distribution of genetic variability, with higher genetic diversity in central populations (e.g., Brazil) and lower diversity in peripheral populations.

**Methods:** Utilizing microsatellite markers, we analyzed larvae collected from infested livestock across South America. Larvae were collected directly from various wound sites to ensure a broad representation of genetic diversity.

**Results:** Contrary to our initial hypothesis, the results revealed consistent genetic variability across the species’ distribution, low population differentiation, and no evidence of isolation-by-distance patterns among subpopulations. The genetic analysis indicated an excess of homozygotes, potentially due to the Wahlund effect, null alleles, or selection pressure.

**Conclusions:** These findings suggest a complex metapopulation structure for *Co. hominivorax*, challenging classical population genetics models. This complexity likely arises from the species’ high dispersal capability and frequent local extinctions followed by recolonization. These results have important implications for the design and implementation of control programs, emphasizing the need for coordinated and large-scale actions rather than isolated initiatives.

## Background

Parasitic diseases significantly influence livestock production, especially in underdeveloped and developing nations [1,2]. These diseases can lead to a range of detrimental effects on livestock, including increased mortality rates, decreased fertility, weight loss, and diminished milk production, thereby causing a significant impact on production levels, both directly and indirectly [3]. Ecto- and endoparasites are the main culprits, recognized for restricting animal production by compromising the health of cattle, sheep, goats, and other livestock species. Given that over a billion people worldwide rely on livestock, managing and controlling parasitic infections has become a high priority to maintain food security and reduce economic losses [4].

Myiasis, an infection caused by Diptera larvae feeding on living or dead tissues of a vertebrate host, liquid body material, or ingested food [5], is a significant concern for livestock production. Several fly species have been identified as responsible for some form of myiasis, with the most important species belonging to the families Muscidae, Calliphoridae, Oestridae, and Sarcophagidae [6]. These species hold medical, sanitary and economic importance, affecting humans and other animals, while contributing to the spread of pathogenic and agricultural losses [5,7].

The New World screwworm (NWS) fly, *Cochliomyia hominivorax* [8] is an obligate ectoparasite of homeothermic animals [5]. Adult females of this species deposit their eggs in wounds or cavities on the host’s body, with the larvae feeding on living tissue. If not treated, wounds can expand, attracting other females to lay their eggs in the same host [9,10]. This species is a major pest in the Neotropical region, infesting domestic animals, mostly cattle [11]. While infestation by screwworm larvae can lead to the host’s death in extreme cases, common manifestations include abortion, reduction in milk production, weight, and fertility [12]. Additionally, infestation scars reduce leather quality, affecting its value [13]. Economic losses caused by this species are substantial, encompassing not only reduced productivity and death, but also the costs of manpower and insecticides involved in the management, prophylaxis, and treatment. Estimates indicate that annual costs in South America can reach 3.6 billion dollars annually [14].

Originally, the geographical distribution of the screwworm fly extended from the southern United States to the northern regions of Argentina and Uruguay [7]. Significant economic losses throughout its distribution led to initiatives to control the fly through male sterilization and the implementation of an eradication program, the Sterile Insect Technique (SIT) [15]. The *Co. hominivorax* eradication program by SIT began in 1958 in the United States, leading to the country being declared free of cases by 1966. Similar successes were achieved throughout Central America to Panama in 2001, with a permanent barrier maintained since then to prevent recolonization by flies from South America [16–19]. Despite these efforts, outbreaks such as the one in Florida Keys in 2016 have occurred but were successfully re-eradicated within six months [20].

Due to the high cost of SIT, the control of *Co. hominivorax* in its current geographical distribution relies on insecticides, mostly organophosphates. However, their extensive use poses serious risks, including toxicity to animals and meat consumers, environmental pollution, and the selection of resistant strains [21–23]*. Cochliomyia hominivorax* is a significant agricultural pest with wide dispersal and adaptability, necessitating continuous investment in monitoring, treatment, and control. Therefore, research in genetics and ecology is crucial for developing effective control strategies. The status of *Co. hominivorax* as a model species for a well-established control program underscores its significance, motivating the study of its biology. Among these, genetic variability and analysis of natural population structure are essential. By investigating these patterns, valuable insights into NWS management can be gained, leading to the development of targeted interventions to mitigate its impact on agricultural systems.

### Population genetics of Co. hominivorax

The first genetic studies for *Co. hominivorax* aimed at characterizing mutations caused by the exposure of pupae to radiation [24]. Almost 20 years later, genetic studies were resumed using isozymes to characterize the variation generated by pupae irradiation. Subsequently, Bush and Neck [25] identified variations in two loci of enzymes related to the production of energy for flight, aGPDH, and PGM, when analyzing samples from the mass rearing of *Co. hominivorax*.

McInnis [26], Richardson et al. [27,28], and Azeredo-Espin [29] initiated studies in North America and Brazil, aiming at the characterization of natural populations with karyotypic, morphological, and sexual compatibility analyses. These studies found a great intra and interpopulation variability in the *Co. hominivorax* populations. Dev et al. [30] used cytological analysis of polytene chromosomes from samples of *Co. hominivorax* from 10 different localities (nine in Mexico and one in Jamaica) but did not identify reproductively isolated populations. In the late 1980s, molecular markers began to be used to describe the genetic variability and structure of natural populations of *Co. hominivorax*.

Initially, mitochondrial DNA (mtDNA) restriction profiles were used to analyze geographic populations of *Co. hominivorax* from the United States, Mexico, Costa Rica, Guatemala, and Jamaica [31,32]. The authors found a high variability, significant population structure, and reduced gene flow, mainly between continental samples and samples from the island of Jamaica.

Krafsur and Whitten [33], using three polymorphic isozyme loci, found no population differentiation in 11 geographic samples from Mexico. They concluded that *Co. hominivorax* constitutes a single panmictic population. A high level of gene flow was also found for *Co. hominivorax* from different locations in Costa Rica, indicating the absence of genetic structuring [34]. Comparing previously analyzed samples from Mexico and Costa Rica with samples collected in Rio de Janeiro and Rio Grande do Sul, the authors obtained concordant results [35]. The results indicated there was no evidence of population structuring, and the hypothesis formulated was that there was a high rate of gene flow between populations.

On the other hand, mtDNA analysis of geographic populations of *Co. hominivorax* from Brazil revealed a high intra and interpopulation mitochondrial genotypes variability [36,37]. Studies with RFLP, RAPD, and isozymes indicated that the analyzed samples of *Co. hominivorax* present high levels of variability and genetic differentiation, indicating population structuring and reduced gene flow [37–39].

In a study carried out with samples from seven locations in Uruguay [40], no population differentiation was detected. This fact is attributed to the absence of geographical barriers in Uruguay and the passive transport of larvae by host transport. In Uruguay, Torres et al. [41] utilized mtDNA and microsatellite markers and did not identify significant population structuring, observing greater variation within the seven populations than between them, and all populations with remarkably similar allele distributions. Lyra et al. [42] identified a moderate structuring of *Co. hominivorax* populations in a study across 34 locations spanning 10 continental and island countries in Central and South America. In this study, analysis of mitochondrial DNA (mtDNA) data using PCR-RFLP technique unveiled a complex pattern of population genetic structure, with an analysis focusing solely on the islands revealed significant population structure (Ф_ST_ = 0.5234; P < 0.001) and low population variability, indicating that the islands function as independent evolutionary units connected by limited gene flow. By contrast, high variability and low, but significant, differentiation was found among mainland populations (Ф_ST_ = 0.0483; P < 0.001), which could not be attributed solely to geographic distance. Torres and Azeredo-Espin [43] utilized 12 microsatellites to investigate the population genetics of *Co. hominivorax* in the Caribbean. They detected moderate population structure among populations (F_ST_ = 0.157) and high population differentiation, suggesting highly structured populations resulting from either limited gene flow or a source-sink dynamic, along with rapid recovery from population contractions.

In an exploration of the phylogeographic history of *Co. hominivorax*, Fresia et al. [44] delved into mitochondrial DNA sequences from 361 individuals sampled across 38 locations within its contemporary range. Their findings revealed significant genetic divergence on a macrogeographic scale (Φ_ST_ = 0.496; P < 0.001), suggesting historical events as primary drivers shaping genetic diversity distribution, such as Caribbean Island colonization, Amazon region vicariance, and population expansion. In a more recent study, Tietjen et al. [45] found a spectrum of population genetic differentiation from “moderate” to “very large” levels employing SNPs to elucidate the population structure of *Co. hominivorax* at 12 sites spanning its entire distribution area.

These previous studies of natural populations of *Co. hominivorax* have yielded different results, which may appear contradictory. However, a discernible pattern for the distribution of genetic variability in this species can be inferred. In general, these results have indicated a low to moderate genetic structuring in the regions situated at the distribution extremes of this species, contrasting with a high structuring observed in central regions. This pattern suggests a center-periphery distribution of genetic variability. The central populations in Brazil, considered the origin center of the species [39], would harbor older populations and display a pattern of historical differentiation. Populations situated at the distribution peripheries would result from recent colonization events and hence exhibit low levels of population structuring. In this scenario, there would be a population structure following the isolation-by-distance model, with peripheral populations presenting a lower genetic variability than those at the center of the distribution. In light of these observations, our study aimed to investigate the genetic variability and assess the degree of genetic structuring among *Co. hominivorax* geographical populations from South America using microsatellite markers, testing the hypothesis of center-periphery distribution of genetic variability.

## Material and Methods

### Obtaining Co. hominivorax samples from South America

*Cochliomyia hominivorax* larvae were collected directly from wounds of infested animals on livestock farms at eight locations in Brazil. Samples from the remaining locations were obtained from wounds of infested animals and shipped in absolute ethanol by local farmers and collaborators (Table 1, Fig. 1). The samples were collected between 2001 and 2006. These samples were the same samples Lyra and colleagues analyzed in 2009, and Fresia and colleagues in 2011, both with mitochondrial markers [42,44].

**Figure 1.**
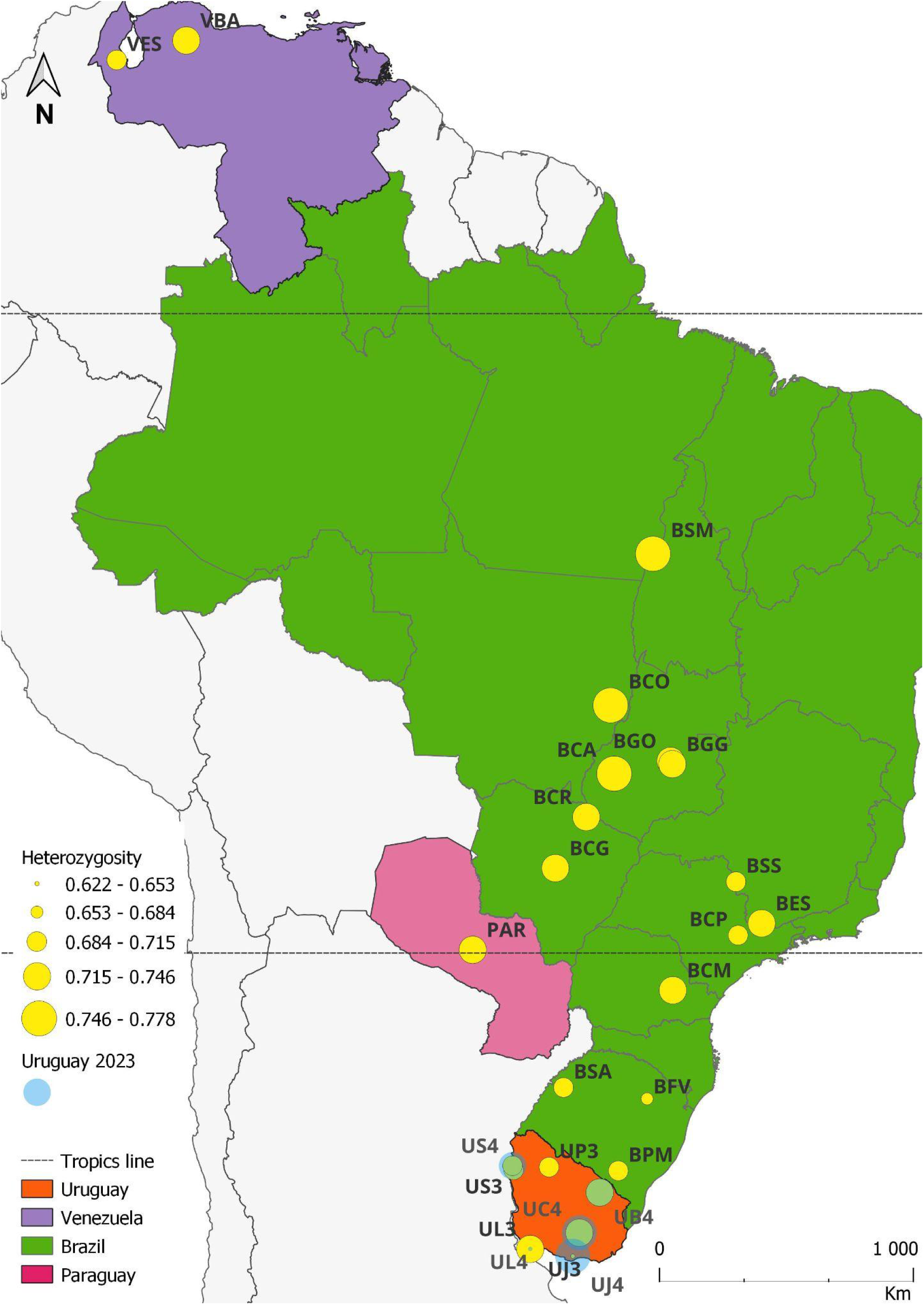
Location of samples of *Co. hominivorax* in South America, with heterozygosity ranging from 0.621 in Joaquín Suarez (UJ3) to 0.778 in Caiapônia (BCA).

**Table 1.**
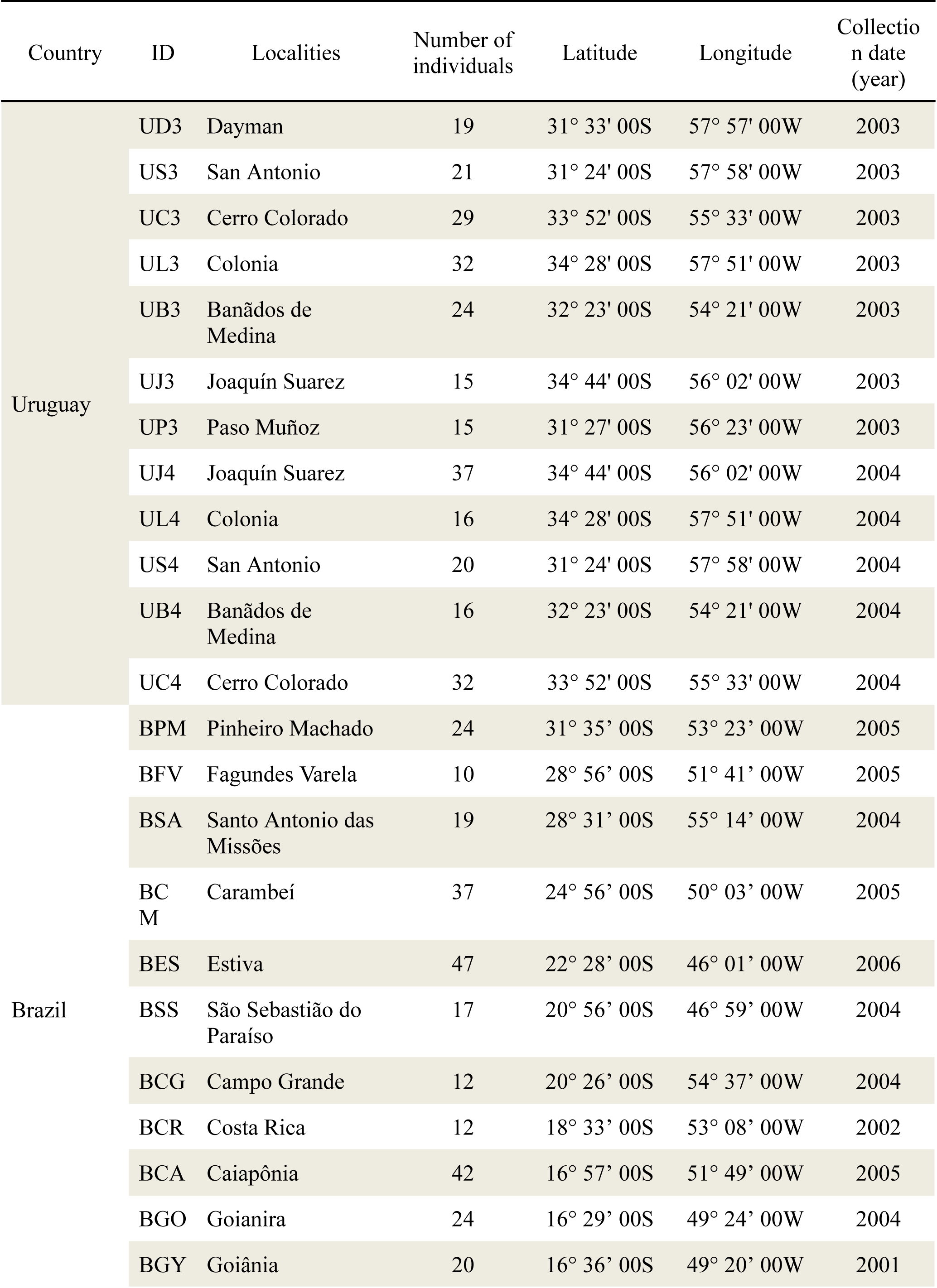
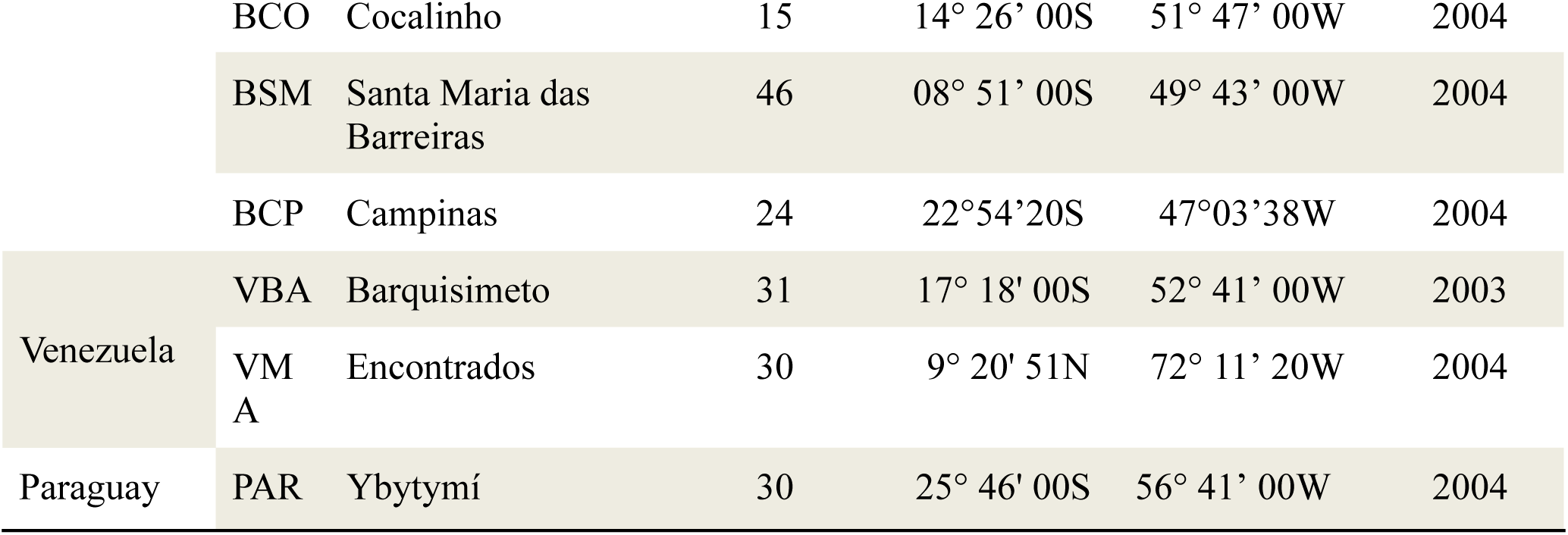
Sampled locations with their respective geographical information.

### Analysis of microsatellite data

#### Detection of microsatellite loci polymorphism

To analyze the geographic populations of South America, we used twelve loci selected from previous work [46,47]. PCR amplification of microsatellite loci detailed in “Additional file 1: Table S1.” and visualization were performed as previously described by Torres et al. ([46]; Table S1).

The number and frequency of alleles, as well as the observed (H_O_) and unbiased expected (H_E_) [48] heterozygosities under Hardy-Weinberg equilibrium, were determined per locus for each location. The ‘basic.stats’ function of the R package hierfstat version 0.5.10 [49] was applied to calculate average observed heterozygosity (H_O_), expected heterozygosity (H_E_), and inbreeding coefficients (F_IS_). Deviations from Hardy-Weinberg equilibrium expectations were assessed for each locus and population using exact tests implemented in GENEPOP 4.7 [50,51].

Allele frequency data were utilized, and statistical tests applied to detect signatures of heterozygosity excess (H_Exc_), indicative of a recent bottleneck event, using the program BOTTLENECK 1.2.02 for all microsatellite loci [52]. The sign test was employed to evaluate the number of loci exhibiting heterozygosity excess compared to the expected number by chance under different mutational models, including the infinite alleles model (IAM), the stepwise mutation model (SMM), and the two-phase model (TPM). For the TPM we chose the proportions in favor of IAM (30% of SMM and 70% of IAM).

Interpopulation differentiation was measured in terms of *F*_ST_ estimates in samples from South America. Three sets of subpopulations were used to calculate interpopulation differentiation in terms of *F*_ST_ estimation, inbreeding coefficient F_IS_ and estimation number of migrants *Nm*. The first set includes all samples from Table 1, excluding only samples obtained from Uruguay in 2004. The second set excludes only samples obtained from Uruguay in 2003. The third group includes all samples from Table 1, comprising both samples from Uruguay in 2004 and 2003 without grouping them. The genetic differentiation index, F_ST_, was calculated between localities using absolute allele frequencies, following the methodology described by Weir and Cockerham [53], implemented via the hierfstat library in R [49]. Pairwise F_ST_ estimates were assessed for significance through genotype permutation among populations, as outlined by Goudet et al. [54]. The critical significance level applied in all statistical tests was 0.05. In all simultaneous statistical tests, critical significance levels were corrected using the Sequential Bonferroni test [55] to enable the overall significance to be examined. The isolation-by-distance model, assessing population genetic structure, was evaluated through linear regression. This involved correlating pairwise F_ST_ /(1 - F_ST_) with the natural logarithm of the geographical distance between population pairs [56]. Additionally, comparisons were made between the sample groups’ genetic variability using “Wilcoxon signed rank” tests. These analyses were performed on the R [57] computational platform.

## Results

### Genetic variability

The number of alleles observed heterozygosity (H_O_), and expected heterozygosity (H_E_) were calculated per locus and population (Additional file 2: Table S2; Additional file 3 Fig. S1). The number of alleles detected per locus ranged from 2 to 18, with an average of 6.9 alleles per locus and per population. The expected heterozygosity ranged from 0.194 (Costa Rica, Brazil, locus CH09) to 0.936 (Costa Rica, Brazil, locus CH21), with an overall average of 0.722. With the exclusion of samples from Uruguay, the average did not change, and it was 0.727. Significant deviation from the Hardy-Weinberg equilibrium were found in 187 out of 348 tests, though after sequential Bonferroni correction, this number reduced to 124 significant tests. Such deviations were consistent across all analyzed samples, manifesting in at least one locus per location. Linkage disequilibrium analysis revealed that 376 comparisons between loci pairs out of 1782 showed linkage disequilibrium (p < 0.05). Additionally, this analysis revealed non-random associations between loci pairs in 56 comparisons After sequential Bonferroni correction. These imbalances varied across subpopulations.

The analysis of all the loci revealed the least variability in expected heterozygosity in Colonia del Sacramento, Uruguay (2004), with H_E_ equal to 0.633 (Fig. 1), significantly different from the overall mean (Wilcoxon Signed-rank test, p = 0.001). Similarly, Joaquín Suarez, Uruguay (2003), exhibited lower mean variability (H_E_ = 0.622; Wilcoxon Signed-rank test, p = 0.0243). Conversely, the highest observed variabilities were recorded in Joaquín Suarez, Uruguay (2004) (0.774; p = 0.02), Santa Maria das Barreiras, Brazil (0.775; p = 0.0018), and Caiapônia, Brazil (0.778; p = 0.0001). Heterozygosity was also significantly different from the overall mean in the populations of San Antonio, Uruguay (2003 and 2004), Cerro Colorado, Uruguay (2004), and Campinas, Brazil, as shown in “Additional file 2: Table S2”. Other observed differences from the mean were not statistically significant.

To test the decrease in the genetic variability in the extreme south of the *Co. hominivorax* distribution, a test was carried out comparing the H_E_ of the southern locations (below the Capricorn Tropic), including all the subpopulations of Uruguay and the four of Southern Brazil with populations of the central region (the remaining Brazil and Venezuela subpopulations). The *Co. hominivorax* variability in the south (H_E_ = 0.7206) was not significantly different from the variability of central populations (H_E_ = 0.7372), with p = 0.251. Additionally, no significant difference was observed when comparing allelic diversities (“allelic richness”, DA) between southern and central groups (DA_south_ = 4.71; DA_central_ = 4.94; p = 0.10987) or when considering only samples from Uruguay as a southern group (H_E Uruguay_ = 0.7206, H_E central_ = 0.7372 and p = 0.5658; DA_Uruguay_ = 4.84, DA_central_ = 4.94 and p = 0.16967).

A bottleneck was not detected in any of the subpopulations across all three mutation models simultaneously (Table 2). However, in most of them, except for San Antonio, Uruguay (2003), Santo Antonio das Missões, and São Sebastião do Paraíso, Brazil, an excess of heterozygotes was detected in relation to the number of heterozygotes expected at equilibrium in the IAM model. Under the SMM model, nine populations putatively experienced a reduction in population size (San Antonio and Banãdos de Medina 2003, Pinheiro Machado, Santo Antonio das Missões, Carambeí, Estiva, São Sebastião do Paraíso, Caiapônia, and Goiânia). Under TPM, there was a significant population reduction in San Antonio (2004) and Cerro Colorado (2004).

**Table 2.**
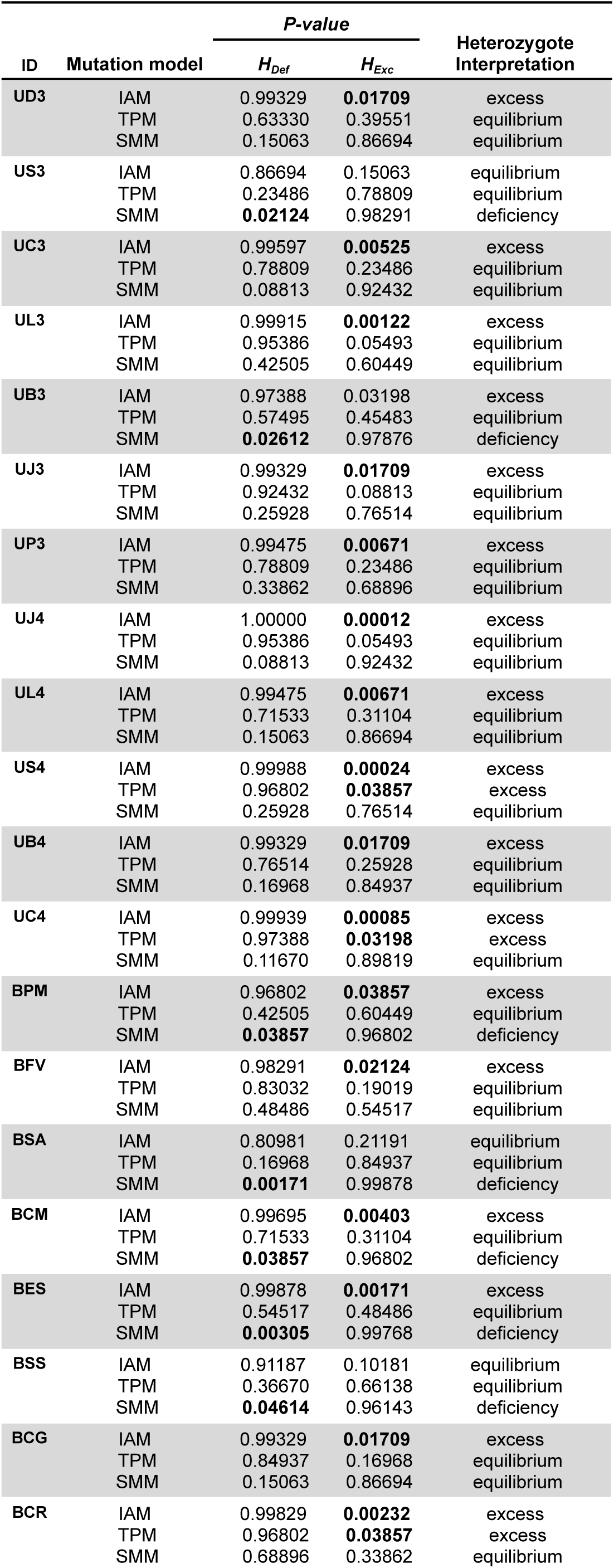
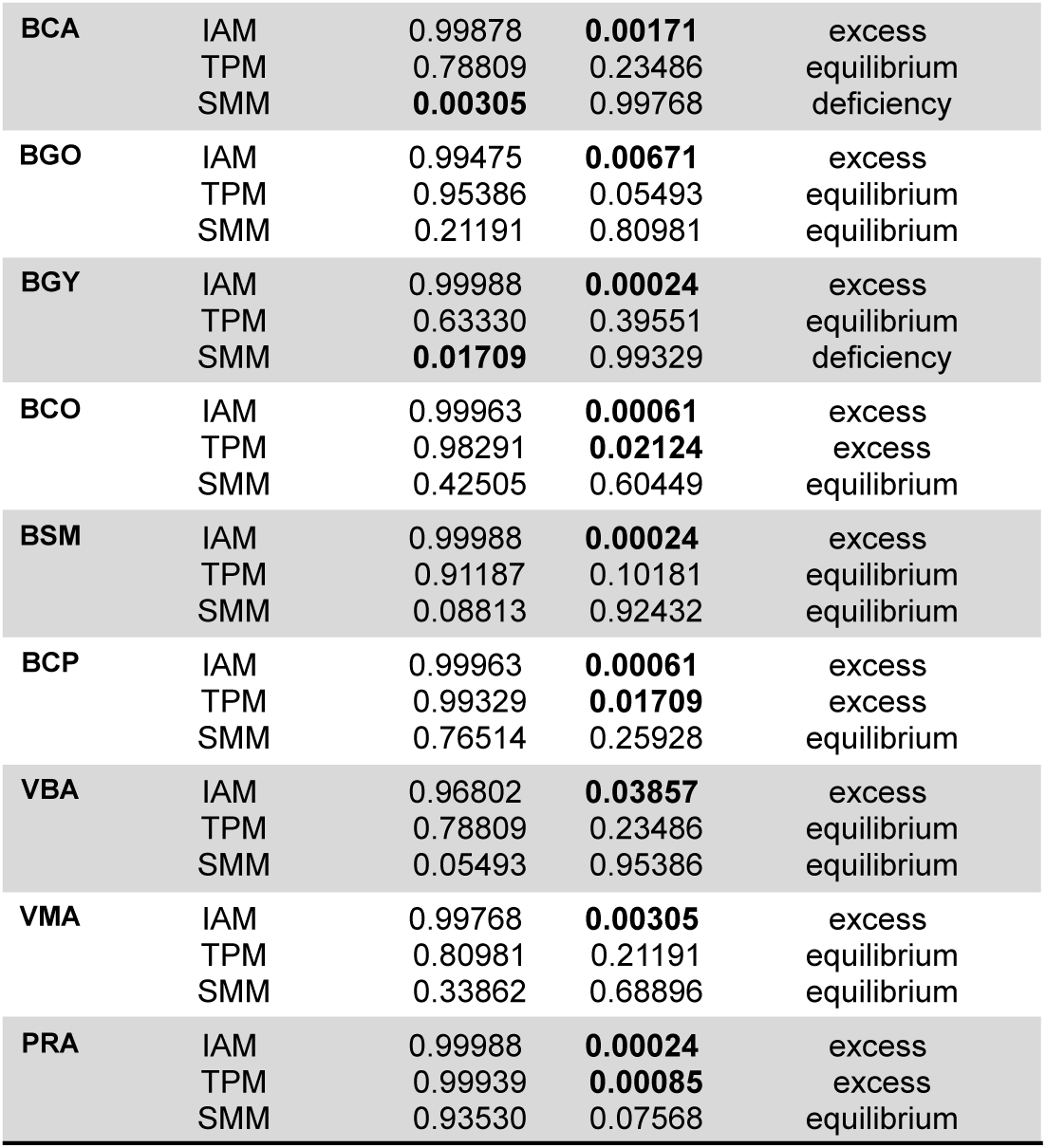
Tests to detect the populational reduction on the BOTTLENECK program.

| ID | Mutation model | *P-value* |  | Heterozygote Interpretation |
| --- | --- | --- | --- | --- |
| | | $H_{Def}$ | $H_{Exc}$ | |
| UD3 | IAM | 0.99329 | **0.01709** | excess |
|  | TPM | 0.63330 | 0.39551 | equilibrium |
|  | SMM | 0.15063 | 0.86694 | equilibrium |
| US3 | IAM | 0.86694 | 0.15063 | equilibrium |
|  | TPM | 0.23486 | 0.78809 | equilibrium |
|  | SMM | **0.02124** | 0.98291 | deficiency |
| UC3 | IAM | 0.99597 | **0.00525** | excess |
|  | TPM | 0.78809 | 0.23486 | equilibrium |
|  | SMM | 0.08813 | 0.92432 | equilibrium |
| UL3 | IAM | 0.99915 | **0.00122** | excess |
|  | TPM | 0.95386 | 0.05493 | equilibrium |
|  | SMM | 0.42505 | 0.60449 | equilibrium |
| UB3 | IAM | 0.97388 | 0.03198 | excess |
|  | TPM | 0.57495 | 0.45483 | equilibrium |
|  | SMM | **0.02612** | 0.97876 | deficiency |
| UJ3 | IAM | 0.99329 | **0.01709** | excess |
|  | TPM | 0.92432 | 0.08813 | equilibrium |
|  | SMM | 0.25928 | 0.76514 | equilibrium |
| UP3 | IAM | 0.99475 | **0.00671** | excess |
|  | TPM | 0.78809 | 0.23486 | equilibrium |
|  | SMM | 0.33862 | 0.68896 | equilibrium |
| UJ4 | IAM | 1.00000 | **0.00012** | excess |
|  | TPM | 0.95386 | 0.05493 | equilibrium |
|  | SMM | 0.08813 | 0.92432 | equilibrium |
| UL4 | IAM | 0.99475 | **0.00671** | excess |
|  | TPM | 0.71533 | 0.31104 | equilibrium |
|  | SMM | 0.15063 | 0.86694 | equilibrium |
| US4 | IAM | 0.99988 | **0.00024** | excess |
|  | TPM | 0.96802 | **0.03857** | excess |
|  | SMM | 0.25928 | 0.76514 | equilibrium |
| UB4 | IAM | 0.99329 | **0.01709** | excess |
|  | TPM | 0.76514 | 0.25928 | equilibrium |
|  | SMM | 0.16968 | 0.84937 | equilibrium |
| UC4 | IAM | 0.99939 | **0.00085** | excess |
|  | TPM | 0.97388 | **0.03198** | excess |
|  | SMM | 0.11670 | 0.89819 | equilibrium |
| BPM | IAM | 0.96802 | **0.03857** | excess |
|  | TPM | 0.42505 | 0.60449 | equilibrium |
|  | SMM | **0.03857** | 0.96802 | deficiency |
| BFV | IAM | 0.98291 | **0.02124** | excess |
|  | TPM | 0.83032 | 0.19019 | equilibrium |
|  | SMM | 0.48486 | 0.54517 | equilibrium |
| BSA | IAM | 0.80981 | 0.21191 | equilibrium |
|  | TPM | 0.16968 | 0.84937 | equilibrium |
|  | SMM | **0.00171** | 0.99878 | deficiency |
| BCM | IAM | 0.99695 | **0.00403** | excess |
|  | TPM | 0.71533 | 0.31104 | equilibrium |
|  | SMM | **0.03857** | 0.96802 | deficiency |
| BES | IAM | 0.99878 | **0.00171** | excess |
|  | TPM | 0.54517 | 0.48486 | equilibrium |
|  | SMM | **0.00305** | 0.99768 | deficiency |
| BSS | IAM | 0.91187 | 0.10181 | equilibrium |
|  | TPM | 0.36670 | 0.66138 | equilibrium |
|  | SMM | **0.04614** | 0.96143 | deficiency |
| BCG | IAM | 0.99329 | **0.01709** | excess |
|  | TPM | 0.84937 | 0.16968 | equilibrium |
|  | SMM | 0.15063 | 0.86694 | equilibrium |
| BCR | IAM | 0.99829 | **0.00232** | excess |
|  | TPM | 0.96802 | **0.03857** | excess |
|  | SMM | 0.68896 | 0.33862 | equilibrium |
| **BCA** | IAM | 0.99878 | **0.00171** | excess |
|  | TPM | 0.78809 | 0.23486 | equilibrium |
|  | SMM | **0.00305** | 0.99768 | deficiency |
| **BGO** | IAM | 0.99475 | **0.00671** | excess |
|  | TPM | 0.95386 | 0.05493 | equilibrium |
|  | SMM | 0.21191 | 0.80981 | equilibrium |
| **BGY** | IAM | 0.99988 | **0.00024** | excess |
|  | TPM | 0.63330 | 0.39551 | equilibrium |
|  | SMM | **0.01709** | 0.99329 | deficiency |
| **BCO** | IAM | 0.99963 | **0.00061** | excess |
|  | TPM | 0.98291 | **0.02124** | excess |
|  | SMM | 0.42505 | 0.60449 | equilibrium |
| **BSM** | IAM | 0.99988 | **0.00024** | excess |
|  | TPM | 0.91187 | 0.10181 | equilibrium |
|  | SMM | 0.08813 | 0.92432 | equilibrium |
| **BCP** | IAM | 0.99963 | **0.00061** | excess |
|  | TPM | 0.99329 | **0.01709** | excess |
|  | SMM | 0.76514 | 0.25928 | equilibrium |
| **VBA** | IAM | 0.96802 | **0.03857** | excess |
|  | TPM | 0.78809 | 0.23486 | equilibrium |
|  | SMM | 0.05493 | 0.95386 | equilibrium |
| **VMA** | IAM | 0.99768 | **0.00305** | excess |
|  | TPM | 0.80981 | 0.21191 | equilibrium |
|  | SMM | 0.33862 | 0.68896 | equilibrium |
| **PRA** | IAM | 0.99988 | **0.00024** | excess |
|  | TPM | 0.99939 | **0.00085** | excess |
|  | SMM | 0.93530 | 0.07568 | equilibrium |

### Genetic differentiation

Genetic differentiation index (F_ST_) were significant and small across subpopulation groups (Table 3), with moderate inbreeding coefficient (Table 3) (F_IS_), and moderate estimated number of migrants (Table 3). The global *F*_ST_ estimates were low but significant, indicating a population structure for the analyzed samples. Even in a species with a high dispersal rate, these values are surprisingly low given the vast territorial extension covered (samples from up to 5000 km away were compared).

The *F*_ST_ pairwise estimates were also generally low and different from zero for 264 out of 276 pairs within group 1 (Fig. 2a), and for 220 out of 231 pairs within group 2 (Fig. 2b). For group 3, the estimates were also different from 0 for 371 out of 406 pairs, ranging from 0.00686 (between Colonia, Uruguay 2003 and Banãdos de Medina, Uruguay 2003) to 0.13815 (between Joaquín Suarez, Uruguay 2003 and Colonia, Uruguay 2004). Nine subpopulations that presented all significant estimates in group 3 (Joaquín Suarez, Uruguay 2004; Colonia, Uruguay 2004; San Antonio Uruguay 2003, 2004; Banãdos de Medina 2004; Cerro Colorado 2003, 2004; Campinas, Brazil; Barquisimeto and Encontrados, Venezuela) differed from all other populations. Nevertheless, estimates were low, ranging from 0.0251 (with São Sebastião do Paraíso) and 0.1193 (with Colonia del Sacramento (2004), Uruguay) (Fig. 2c).

**Figure 2.**
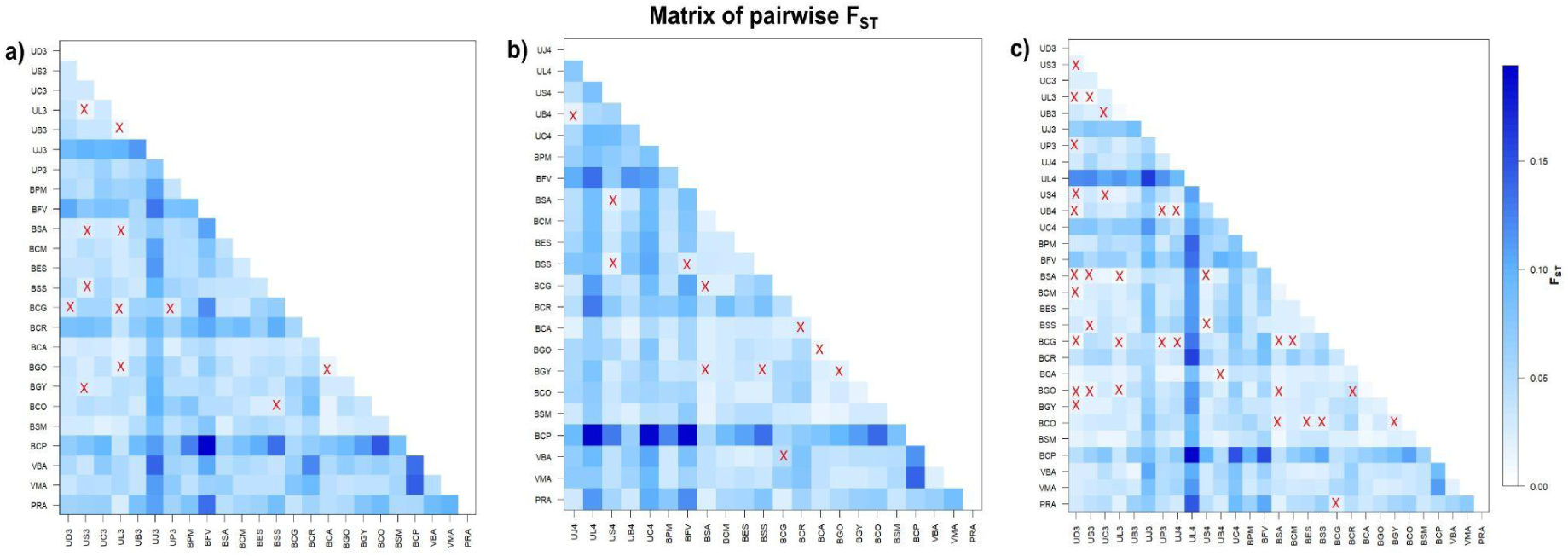
Matrix of pairwise FST estimates between subpopulations pairs a) Group 1 b) Group 2, c) Group 3, “x”, not statistically significant FST pairwise.

The correlation between the genetic and geographic distances was not significant after the Mantel test neither in the group 1 samples (p = 0.1954, Fig. 3a) nor in the group 2 samples (p = 0.137. Fig. 3b). This pattern did not change even when excluding locations separated by distances inferior to 100 km (group 1, p = 0.116; group 2, p = 0.158) or 200 km (group 1, p = 0.134; group 2, p = 0.152). Tests using other minimum and maximum distances also failed to indicate any genetic and geographic distance association (data not shown).

**Figure 3.**
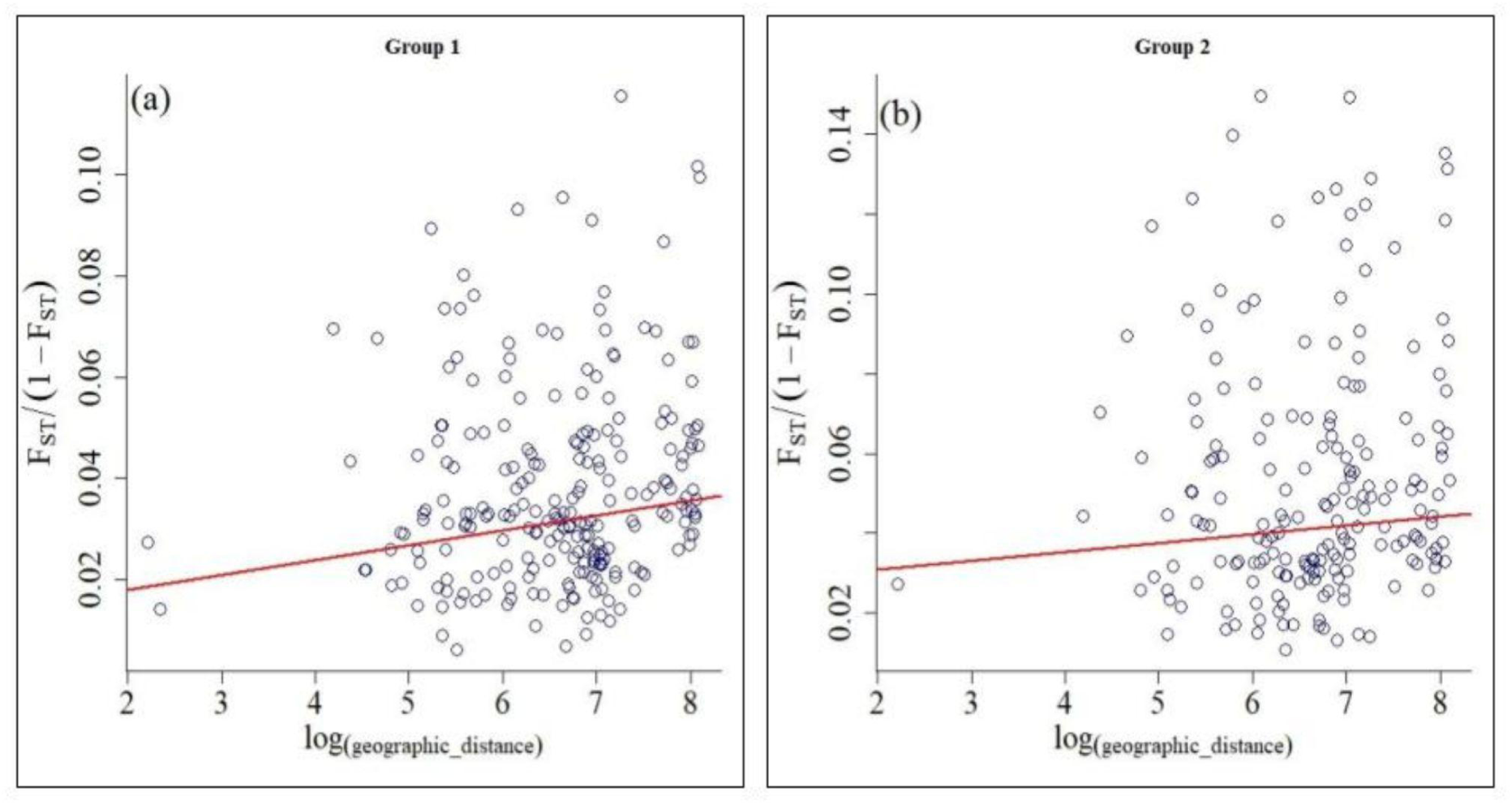
Test for isolation by distance (IBD). Linear regression of *F_ST_* /(1-*F_ST_*) between population pairs against the natural logarithm of geographical distances between population pairs, both not significant. (a) group 1 subpopulation; (b) group 2 subpopulation.

The low differentiation found between subpopulations was not expected at such large distances (for example, F_ST_ = 0.0534 estimation between samples presenting the greatest geographic distance, 5180 km).

We observed temporal substructuring of *Co. hominivorax* subpopulations in the southern region of Uruguay, specifically in Joaquín Suarez and Colonia. These populations exhibit significant differences in heterozygosity within the same locations across different years (H_E_ for UJ3 = 0.66 and UJ4 = 0.82; UL3 = 0.77 and UL4 = 0.63), with greater differentiation (F_ST_ pairwise) compared to other locations (F_ST_ pairwise UJ3 and UJ4 = 0.074; UL3 and UL4 = 0.095).

## Discussion

According to our initial hypothesis, based on previous studies on *Co. hominivorax* populations, we anticipated a high genetic differentiation in subpopulations with a central distribution, structured according to an isolation-by-distance model. Additionally, peripheral populations were expected to exhibit reduced genetic variability compared to central populations. However, we found that: (1) genetic variability remains consistent across the species’ distribution extremes; (2) population differentiation is low, and (3) subpopulations do not follow an isolation-by-distance model; (4) temporal substructuring in peripheral populations.

Our results contrasted from those obtained by Infante-Vargas and Azeredo-Espin [38], and Infante-Malachias [37]. Using mtDNA, RFLP markers, RAPD, and Isozymes, they indicated high levels of genetic differentiation with reduced gene flow among Brazilian *Co. hominivorax* populations Nevertheless, Lyra et al. [42] found lower but significant differentiation among mainland populations using mtDNA polymerase chain reaction-restricted fragment length polymorphism (PCR-RFLP; ф_ST_ = 0.039, p < 0.05). Discrepancies among these studies might be due to differences in molecular markers and sampled locations.

The main difference between our study and the previous ones lies in the sampling methodology. Due to difficulties in using artificial attractants to collect *Co. hominivorax* adults in traps, Lyra et al. [42] and we collected larvae directly from infested animals, ensuring a wider sampling collection. However, this approach may generate biases if individuals from a single oviposition are overrepresented. This bias might occur due to larvae from a single open wound are from an oviposition by only one or a few females [9]. If all individuals from a wound are collected, the sample size may be adequate at 20-30 individuals, but they may be from one or two families, which would bias the results. In previous studies, between 16 and 27 individuals were used per wound. Individuals were sampled from one, two, three, or nine wounds in each analyzed subpopulation. The use of many individuals per wound associated with sampling in only a few wounds may have generated deviations due to the increase in the frequency of rare haplotypes or alleles and the failure to observe all possible haplotypes or genotypes in each locus. To mitigate this, we implemented the criteria for selecting larvae from different ovipositions, broadening our sampling across wounds from different hosts. In addition, larval stage and mitochondrial haplotype from previous studies [40,42] were used to differentiate individuals that were on the same wound but came from different ovipositions.

In the current analysis using all loci together, all samples exhibited an excess of homozygotes compared to the expected equilibrium, potentially attributable to different factors, including the Wahlund effect, null alleles, and selection pressure [58]. Firstly, these deviations can stem from the Wahlund effect [59], which arises when a population is not homogeneous but consists of subpopulations with varying allele frequencies [60]. When samples from these subpopulations are mixed, it can lead to an excess of homozygotes compared to what would be expected if all individuals were mating randomly [61,62]. This was confirmed by the positive correlation observed between F_IS_ and F_ST_.

The excess of homozygotes could also be due to the presence of null alleles, which might lead to an underestimation of heterozygosity [63]. Although null alleles may be present at some loci, this does not seem to be the only one responsible for the excess of homozygotes since the deviations were generally observed in at least one locus in all subpopulations. Furthermore, there was no change in these results with the exclusion of the locus with the highest null alleles estimate (CH10, data not shown). Once again, the occurrence of demographic changes appears as a possible explanation for the results.

Lastly, the excess of homozygotes may be due to selection acting on one or more loci, if one or more loci are under selection pressure, it can lead to an excess of homozygotes if certain alleles are being favored over others [64]. The control of *Co. hominivorax* with insecticides has caused rapid selection for resistance in some populations [21–23]. This resistance is multilocus, potentially affecting the entire genome and influencing a wide range of genetic traits and interactions [23]. Baqir and Ab Majid [65] highlight selection pressure in the population genetic structure of tropical bugs in Iraq. Their study reveals bottleneck events with an excess of homozygosity, suggesting a decrease in the effective size of the bug population due to control activities and the speed at which the population recovered, emphasizing the role of selection in these populations [65].

Further investigation is needed to elucidate the underlying cause of the observed excess of homozygotes. Understanding this phenomenon is crucial for accurately interpreting population genetic dynamics. Therefore, our interpretation of *Co. hominivorax* population structure and dynamics focus on two main areas: (A) historical processes and (B) contemporary demographic processes.

### A. Historical Processes

Estimates of gene flow between species populations are generally based on differentiation indices such as *F_ST_* estimates [66,67]. This approach is based on a few assumptions. The most crucial one, and the one most likely to be violated in many natural populations, is the balance between mutation, migration, and drift. Violation of balance is critical in pest species (organisms that depend on humans and their resources as causative agents of human disease, domestic animals, and cultivated plants), whose effective population sizes are generally high and whose demographic history is recent [68–70].

Lehmann et al. [71] conducted a study of the distribution of genetic variability of isozyme and microsatellite loci among populations of *An. gambiae* from Kenya and Senegal, Africa. These authors found low population differentiation (*F_ST_* = 0.016) and estimated a high level of gene flow between populations (*Nm* > 7.7) separated by more than 6000 km. Besansky et al. [72], analyzing sequence data from a mitochondrial region ND5 gene (665 bp) from seven villages in Kenya and three in Senegal, found homogeneity between the subpopulations of the two countries (*F_ST_* = 0.085). Consequently, the estimates of gene flow were very high (*Nm* = 5.4). Similar results were obtained for *Anopheles arabiensis*, whose populations separated by 7000 km were also homogeneous (*F_ST_* = 0.044, *Nm* = 10.8).

Donelly et al. [68] analyzed the genetic variability of 18 microsatellite loci in African populations of *An. arabiensis* and *An. gambiae*. They verified that the analyzed populations were not in mutation-migration-drift equilibrium. This observation was consistent with the detection in previous studies of low levels of genetic differentiation across the distribution of these two species. Similar patterns were observed in *Ce. capitata* [73], *Ceratitis rosa* and *Ceratitis flavicentris* [74], in *Drosophila melanogaster* [75] and in populations of dung beetles of the species *Aphodius fossor* that uses bovine manure to feed the larvae, demonstrating a high dependence on livestock [76], such as *Co. hominivorax*.

A parallel can be drawn between these studies and the present analysis of *Co. hominivorax* populations. In this study, a deviation from the Hardy-Weinberg equilibrium was observed in almost every test performed. Likewise, a high number of significant linkage disequilibrium tests were observed. These results are accompanied by a low genetic structure in terms of *F_ST_* estimates and the absence of isolation by distance. For *Co. hominivorax*, these results could also be explained by a recent population expansion. This increase probably accompanied the introduction of cattle and livestock in South America after the beginning of the colonization of the continent.

The first record of the introduction of cattle in South America is from 1524 in the present-day Colombia [77], and livestock is registered as standard practice from 1614 when it became a population factor in the countryside region. Two scenarios can be considered regarding the *Co. hominivorax* population expansion in South America. In the first one, the species was already found in the continent before the introduction of cattle, living in small effective populations, infesting small and medium-sized wild animals. The introduction of intensive livestock farming represented the creation of a new niche for *Co. hominivorax*, causing rapid and disorganized population growth. In the second scenario, this species was not native to the continent, but it was introduced, becoming a problem for livestock in South American countries.

The *Cochliomyia* genus is formed by four species, *Co. hominivorax* (Coquerel), *Cochliomyia macellaria* (Fabricius), *Cochliomyia aldrichi* (Del Ponte), and *Cochliomyia minima* (Shanon). Of these species, only *Co. hominivorax* is described as an obligate ectoparasite [78]. The distribution of *Co. macellaria* is sympatric to *Co. hominivorax*. *Cochliomyia aldrichi* is present in Florida (United States), Bermuda, Bahamas, Cuba, Puerto Rico, San Salvador, the Virgin Islands, and the Cayman Islands [78]. There are records of this species also in the Dominican Republic and Antilles [7]. The distribution of *Co. minima* is more restricted, including Jamaica, Cuba, Dominican Republic, Puerto Rico, the Virgin Islands, West Indies, and Florida [7,78]. With these observations, the radiation of the genus could have occurred outside the South American continent, and the two species, *Co. hominivorax,* and *Co. macellaria* could have been introduced in South America. However, previous studies point to South America as the probable center of diversity and origin of the screwworm fly [37]. Although previous studies have comprehensively evaluated the dynamics and genetic structure of populations of this species outside of South America [43], it is still premature to assert the status of *Co. hominivorax* as an introduced species.

Regardless of which scenario best describes the evolutionary history of this species, the human influence on the population structure of this livestock pest and its potential for dispersion and adaptation is evident. Therefore, *Co. hominivorax* should be considered a high-risk species when introduced into a new environment.

Generally, population genetics models ignore the complexity of biological systems. Evolutionary estimates parameters, such as gene flow, are rarely valid because the parameters that define these models are spatially and temporally variable and are very sensitive to evolutionary forces. Conventional models assume that population structure and demographic parameters such as population size and dispersion rates are uniform and constant in time and space. Demographic and genetic balance assumptions are unrealistic and violated, as verified in *Co. hominivorax* populations in South America. *Cochliomyia hominivorax* population structure does not follow classical population genetics models.

Our data reveal a new scenario much more complex than what had been imagined for *Co. hominivorax*. Thus, the implications for the establishment of control programs are not intuitive. Each of the new hypotheses has distinct implications for control programs.

The recolonization after population reduction in the adverse season is accompanied by a rapid *Co. hominivorax* population growth rate that may have ensured the maintenance of genetic variability. This is consistent with the species recovery capacity that suffers a drastic population reduction in the dry season and recovers quickly in the rainy season. This seasonality in the screwworm fly infestation cases is observed throughout its distribution. Another common factor among the analyzed locations is the indiscriminate use of insecticides, which would also cause this fly’s population reduction.

No previous studies were carried out to investigate the dynamics and genetic structure of *Co. hominivorax* populations prior to control programs using SIT. Therefore, it is difficult to predict the outcome of similar programs in South America or the Caribbean islands. If the scenario proposed in this study is confirmed by additional investigations, it is likely that the best strategy for controlling *Co. hominivorax* populations is a synchronized action with a wide geographic range and not the way it is currently carried out through isolated initiatives. It is also vital for the success of the control program to use an integrated system, with the coordinated use of different control methods and the use of climatic restrictions to help reduce the population. Only this way, we can ensure that the refugees will not serve as a source for new recolonizations, which would compromise the effectiveness of the eradication initiative.

Here we provide essential data on the genetic population structure for this species that, together with information obtained from ecological studies, can be used to assess the necessary size of the management unit and the program implementation area, providing fundamental information for decision-making regarding the eradication of this important pest of livestock throughout its current geographic range.

### B. Contemporary Demographic Processes

Analyses of South American samples’ yielded results similar to those presented in the Uruguayan samples [41]. The migration of adult flies between different subpopulations is not enough to explain the low population differentiation since this would result in a positive correlation between geographic and genetic distances.

Passive migration of larvae through infested animals could explain, in part, the observations at a local scale. The transport of animals can occur between different farms of the same owner or in the early stages of infestation (such as eggs or first instar larvae) with animal trade. However, passive migration would not cause the observed patterns at the analyzed scale for several reasons, (1) the cost of transporting animals over long distances is very high, restricting this practice; (2) infestation in more advanced stages is easy to detect, which makes it challenging to trade infested animals; and (3) inspection is more restricted at international borders.

Fountain et al. [69] used 21 microsatellite markers to investigate human-facilitated metapopulation dynamics in *Cimex lectularius*, an emerging pest. They emphasize the impact of colonizer numbers and demographic history on genetic diversity distribution. Pest control leads to local extinctions, while human-facilitated dispersal fosters colonization, shaping metapopulation dynamics. Founder events reduce diversity and increase genetic drift, causing rapid population divergence. Despite low diversity within infestations, genetic differentiation among infestations (F_ST_ = 0.59) highlights a high population structure. Colonization patterns align with the propagule reservoir model, providing insights for bed bug infestation control.

While the acknowledgment of time’s influence on the genetics of animal populations has expanded through the inclusion of time as a variable in research and analysis [79–81], our endeavor represents a pioneering step in integrating the concept of ‘isolation by time’ into genetic analyses of *Co. hominivorax* populations. The results obtained here with the temporal analysis, comparing samples in Uruguay at the same location in two consecutive years, indicate that the local population dynamics of the screwworm fly include population fluctuations dependent on the occurrence of migration, extinction, and recolonization. Temporal genetic differentiation can occur when local extinction occurs, due to significant fluctuations in population size and subsequent colonization of the empty habitat by individuals from populations from more distant locations. Thus, the data obtained fit into a metapopulation model. At the population level, barriers to dispersal and regional selection play a crucial role in shaping the metapopulation configuration, thereby influencing evolutionary dynamics [82].

A metapopulation can be defined as a set of subpopulations of a given species, with each subpopulation occupying a different fragment of a subdivided habitat. Most species are naturally distributed as metapopulations [83,84]: many insects live in trees or other plants, which are a fragment of habitat; many fish and marine invertebrates live on coral reefs; most parasites live on hosts that are effectively fragments of habitat. In the specific case of *Co. hominivorax*, there is a fragmentation of habitats in several aspects: each farm animal where the larvae develop can be considered a fragment, as well as each livestock region, that usually are not continuous, with areas of agricultural production that could act as a form of fragmentation of habitat. Finally, the distribution of forests where there are sites of aggregation of adult flies [85] is currently fragmented due to human action. Thus, according to species distribution in fragmented habitats and the results demonstrated here, each subpopulation of *Co. hominivorax* should be seen as an integral part of a metapopulation. For other species sharing features with *Co. hominivorax*, such as wide distribution and pest status, population structures have already been described using the metapopulation model. Evidence indicates that populations of the blowfly *Phormia regina* exhibit pronounced temporal structure that seems to correspond with seasonal shifts, suggesting that *P. regina* likely operates within a metapopulation framework. Owings et al. [81] stress the significance of spatiotemporal sampling in uncovering the population genetic structure in blowflies, alongside the impact of abiotic variables on these patterns. Their study delved into the temporal population genetic structure of nine populations of *P. regina* over three years, analyzing them at six polymorphic microsatellite loci. The findings suggest a robust temporal structure mirroring seasonal variations, indicative of a metapopulation dynamic. Molecular variance analysis of these populations corroborated significant temporal genetic differentiation, while further analyses revealed correlations between abiotic factors like temperature, humidity, precipitation, and wind speed with the observed genetic subdivisions.

This population dynamics model has also been described for the Mediterranean fly, *Ceratitis capitata*. In an ecological study on a 3000 km^2^ expanse of crops. Israely et al. [86] analyzed spatial and temporal components influencing the distribution pattern of the fly in Israel. In Israel, the fly recovers quickly, even with routine control and a temporary reduction in population density. As well as *Co. hominivorax*, *C. capitata* does not resist the Israeli winter but recolonizes this region during the favorable season from more beneficial areas in the Mediterranean coastal areas and the Jordan River valley. The results obtained in the temporal analysis indicated that in two of the three analyzed locations, the population recovery in early summer would probably be a result of recolonization from other populations and not the growth of a residual population that survived the winter. The conclusion was that fly migration over long distances played a key role in maintaining Mediterranean fly populations in the country. Ke et al. [87] investigated the population dynamics and migration of the *Plutella xylostella* pest in overwintering regions in southern China and Southeast Asia. They conducted samplings over two consecutive years and constructed a population network to analyze contemporary gene flow. They discovered two distinct swarms, with one replacing the other over time, and estimated an average migration distance of about 1000 km, indicating large-scale migration. They observed greater migration activity in spring than in winter, identifying source and sink regions. This rapid population turnover and metapopulation dynamics highlight the importance of temporal sampling and network analysis in understanding contemporary genetic patterns and effectively managing agricultural insect pests like P. *xylostella*. The findings contribute to monitoring insecticide resistance and identifying key populations for long-term sustainable management.

Another example comes from *Anopheles gambiae*, the main insect vector of *Plasmodium falciparum*, the parasite that causes malaria. The genetic diversity of populations of this mosquito is affected by fluctuations in temporal and geographic genetic structure. *An. gambiae* populations are characterized by geographic heterogeneity, absence of random crossing, and absence of limits for dispersion. Microgreographical studies in a village in Mali indicated a metapopulation structure [88]. However, two different metapopulation structures are proposed for this species to try to explain the maintenance of local populations. In the first scenario, local populations disappear entirely during the dry season and are re-established in the favorable season. In the second, local populations are maintained in situ by some form of individual aestivation or diapause in the dry season. With the return of favorable conditions, the individuals in diapause would restore the population. With the data obtained for this species so far, it has not yet been possible to determine which factor is most important in the observed pattern of genetic structure.

In a grasshopper species, *Schistocerca gregaria* [89], a metapopulation structure highly dependent on extinction and recolonization processes was described. This species, together with species of the genus *Locusta*, represents the major pests of agriculture in desert areas of the old world. During the recession period, this species is reduced to small, barely detectable population sizes subject to local extinctions. However, during plague outbreaks, there was a surprising increase in population density, creating grasshoppers’ clouds that attacked crops in as many as 60 countries in the old world. According to the author, the metapopulation structure, mainly the recolonization of unoccupied niches, is fundamental for maintaining the cycles of outbreaks and population retraction of this species. Similarly demonstrated in populations of *Aedes aegypti* from Brazil, Mendonça et al. [90] examined the genetic structure and gene flow of *Ae. aegypti* populations in Manaus, Brazil, during periods of high infestation and dengue outbreaks in urban neighborhoods, both during rainy and dry seasons, using nine microsatellite loci. The results showed genetic homogeneity and extensive gene flow during the rainy season, attributed to abundant breeding sites. Conversely, the dry season exhibited significant genetic structure, primarily due to reduced effective population size in some neighborhoods. Genetic bottleneck analyses indicated continuous population maintenance with seasonal reductions rather than severe bottlenecks. These findings are crucial for the development of effective dengue control strategies.

## Conclusions

The effective size of the *Co. hominivorax* metapopulation in South America is large and dynamic, ensuring theo preservation of genetic diversity and reducing the risk of local extinction due to processes such as inbreeding. The metapopulation has a high probability of survival due to gene flow and recolonization, with migration between subpopulations maintaining overall stability. This stability persists even if some subpopulations undergo local extinctions during cold and dry periods or due to local pesticide use. Although the temporal analysis included only a few populations over a short time period, it was sufficient to demonstrate that populations sampled at different times are distinct. This novel finding, supported by the metapopulation structure of the species, indicates that control strategies should be coordinated and comprehensive. Effective and sustainable pest control must consider the connectivity between subpopulations to optimize management efforts.

## Acknowledgements

Authors are grateful to Rosangela A. Rodrigues, and Maria Salete Couto for their assistance in the laboratory and with fly maintenance.

## Availability of data and materials

All data provided and analysed during this study are included in this article.

## Authors’ contributions

AA-E and TT designed the experiments, KSS and TT analysed the data KS, LC and TT helped draft the manuscript.

## Supplementary information

*Additional file 1: Table S1.* Microsatellite loci polymorphism.

*Additional file 2: Table S2.* Genetic diversity of *Co. hominivorax* samples collected in different locations in South America.

*Additional file 3: Fig S1.* Number of alleles.

## References

1. Fitzpatrick JL. Global food security: The impact of veterinary parasites and parasitologists. Veterinary Parasitology 2013; 195:233–48.

2. Akash, Hoque M, Mondal S, Adusumilli S. Sustainable livestock production and food security [Internet]. In: Emerging Issues in Climate Smart Livestock Production. Elsevier; 2022. page 71–90. Available from: https://linkinghub.elsevier.com/retrieve/pii/B9780128222652000119

3. Kappes A, Tozooneyi T, Shakil G, Railey AF, McIntyre KM, Mayberry DE, et al. Livestock health and disease economics: a scoping review of selected literature. Front. Vet. Sci. 2023; 10:1168649.

4. Sargison N. The critical importance of planned small ruminant livestock health and production in addressing global challenges surrounding food production and poverty alleviation. New Zealand Veterinary Journal 2020; 68:136–44.

5. Zumpt FO. Myiasis in Man and Animals in the Old World. London, Butterworths; 1965.

6. Scholl PJ, Colwell DD, Cepeda-Palacios R. Myiasis (Muscoidea, Oestroidea) [Internet]. In: Medical and Veterinary Entomology. Elsevier; 2019. page 383–419. Available from: https://linkinghub.elsevier.com/retrieve/pii/B9780128140437000194

7. Hall M, Wall R. Myiasis of Humans and Domestic Animals [Internet]. In: Advances in Parasitology. Elsevier; 1995 [cited 2021 Sep 23]. page 257–334. Available from: https://linkinghub.elsevier.com/retrieve/pii/S0065308X08600731

8. Coquerel C. Note sur les larves appartenant a une espèce nouvelle de Diptère (Lucilia hominivorax) developpees dans les sinus frontaux de l’homme a Cayenne. Annales de la Société Entomologique de France 1858; 3:171–6.

9. Thomas DB, Mangan RL. Oviposition and Wound-Visiting Behavior of the Screwworm Fly, Cochliomyia hominivorax (Diptera: Calliphoridae). Annals of the Entomological Society of America 1989; 82:526–34.

10. Wainwright SH, Cunha CW, Webb B, McGregor B, Drolet B, Welch JB. Reemerging/Notifiable Diseases to Watch. Veterinary Clinics of North America: Food Animal Practice 2024; S0749072024000094.

11. Scott MJ, Benoit JB, Davis RJ, Bailey ST, Varga V, Martinson EO, et al. Genomic analyses of a livestock pest, the New World screwworm, find potential targets for genetic control programs. Commun Biol 2020; 3:424.

12. Foerster N, Soresini G, Paiva F, Silva FAD, Leuchtenberger C, Mourão G. First report of myiasis caused by Cochliomyia hominivorax in free-ranging giant otter (Pteronura brasiliensis). Rev. Bras. Parasitol. Vet. 2022; 31:e009522.

13. Lopes LB, Nicolino R, Capanema RO, Oliveira CSF, Haddad JPA, Eckstein C. Economic impacts of parasitic diseases in cattle. CABI Reviews 2016; 1–10.

14. Vargas-Terán M, Spradbery JP, Hofmann HC, Tweddle NE. Impact of Screwworm Eradication Programmes Using the Sterile Insect Technique [Internet]. In: Sterile Insect Technique. Boca Raton: CRC Press; 2021 [cited 2024 Apr 26]. page 949–78. Available from: https://www.taylorfrancis.com/books/9781003035572/chapters/10.1201/9781003035572-29

15. Vreysen MJB, Abd-Alla AMM, Bourtzis K, Bouyer J, Caceres C, De Beer C, et al. The Insect Pest Control Laboratory of the Joint FAO/IAEA Programme: Ten Years (2010–2020) of Research and Development, Achievements and Challenges in Support of the Sterile Insect Technique. Insects 2021; 12:346.

16. Krafsur ES. Sterile Insect Technique for Suppressing and Eradicating Insect Population: 55 Years and Countingl. 1998;15.

17. Wyss JH. Screwworm Eradication in the Americas. Annals of the New York Academy of Sciences 2000; 916:186–93.

18. Baumhover AH. A personal account of developing the sterile insect technique to eradicate the screwworm from curacao, Florida and the southeastern United States. Florida Entomologist 2002; 85:666–73.

19. Klassen W, Curtis CF. History of the Sterile Insect Technique [Internet]. In: Dyck VA, Hendrichs J, Robinson AS, editors. Sterile Insect Technique. Berlin/Heidelberg: Springer-Verlag; 2005 [cited 2023 Mar 23]. page 3–36. Available from: http://link.springer.com/10.1007/1-4020-4051-2_1

20. Skoda SR, Phillips PL, Welch JB. Screwworm (Diptera: Calliphoridae) in the United States: Response to and Elimination of the 2016–2017 Outbreak in Florida. Journal of Medical Entomology 2018; 55:777–86.

21. De Carvalho RA, Torres TT, De Azeredo-Espin AML. A survey of mutations in the Cochliomyia hominivorax (Diptera: Calliphoridae) esterase E3 gene associated with organophosphate resistance and the molecular identification of mutant alleles. Veterinary Parasitology 2006; 140:344–51.

22. Carvalho RA, Torres TT, Paniago MG, Azeredo-Espin AML. Molecular characterization of esterase E3 gene associated with organophosphorus insecticide resistance in the New World screwworm fly, *Cochliomyia hominivorax*. Medical and Veterinary Entomology 2009; 23:86–91.

23. Tandonnet S, Cardoso GA, Mariano-Martins P, Monfardini RD, Cunha VAS, de Carvalho RA, et al. Molecular basis of resistance to organophosphate insecticides in the New World screw-worm fly. Parasites Vectors 2020; 13:562.

24. Kauffman G, Wasserman M. Effects of irradiation on the screwworm, *Callitroga hominivorax* (Cog). University of Texas 1957; 246–59.

25. Bush GL, Neck RW. Ecological Genetics of the Screwworm Fly,Cochliomyia hominivorax(Diptera: Calliphoridae) and Its Bearing on the Quality Control of Mass-reared Insects1. Environmental Entomology 1976; 5:821–6.

26. McInnis DO. Cytogenetics of a Local Population of the Screwworm, Cochliomyia hominivorax,1 from Northeastern Mexico. Annals of the Entomological Society of America 1981; 74:582–9.

27. Richardson RH, Ellison JR, Averhoff WW. Autocidal Control of Screwworms in North America. Science 1982; 215:361–70.

28. Richardson RH, Ellison JR, Averhoff WW. *Response*: Mating Types in Screwworm Populations? Science 1982; 218:1143–5.

29. Azeredo-Espin AML. Análise Cariotípica, Morfométrica e de Compatibilidade Sexual, em Linhagens Brasileiras de Cochliomyia hominivorax (Diptera: Calliphoridae). 1987.

30. Dev V, LaChance LE, Whitten CJ. Polytene chromosomes, karyotype correlations, and population cytology of the primary screwworm fly. Journal of Heredity 1986; 77:427–34.

31. Roehrdanz RL. Intraspecific genetic variability in mitochondrial DNA of the screwworm fly (Cochliomyia hominivorax). Biochem Genet 1989; 27:551–69.

32. Roehrdanz RL, Johnson DA. Mitochondrial DNA Variation among Geographical Populations of the Screwworm Fly, Cochliomyia hominivorax1. Journal of Medical Entomology 1988; 25:136–41.

33. Krafsur ES, Whitten CJ. Breeding Structure of Screwworm Fly Populations (Diptera: Calliphoridae) in Colima, Mexico. Journal of Medical Entomology 1993; 30:477–80.

34. Taylor DB, Peterson RD. Population Genetics and Gene Variation in Primary and Secondary Screwworm (Diptera: Calliphoridae). Ann. Entomol. Soc. Am. 1994; 87:626–33.

35. Taylor DB, Peterson RD, Moya-Borja GE. Population genetics and gene variation in screwworms (Diptera: Calliphoridae) from Brazil. Biochem Genet 1996; 34:67–76.

36. Azeredo-Espin AML. Mitochondrial DNA variability in geographical populations of the brazilian screwworm fly. In: International Atomic Energy Agency, Food and Agriculture Organization of the United Nations, editors. Management of insect pests nuglear and related molecular and genetic techniques. Vienna: [Lanham, MD: The Agency; UNIPUB, Distributor in the U.S.A. and Canada]; 1993. page 161–5.

37. Infante-Malachias ME. Estrutura genetica de populações de Cochliomyia hominivorax (diptera: calliphoridae) da região sudeste do Brasil: analise atraves de tres tipos de marcadores geneticos [Internet]. 1999 [cited 2023 Mar 23]; Available from: http://acervus.unicamp.br/index.asp?codigo_sophia=331435

38. Infante Vargas ME, Azeredo-Espin AML. Genetic VAriability in Mitochondrial DNA of the Screwworm, *Cochliomyia hominivorax* (Diptera: Calliphoridae), from Brazil. Biochem. Genet. 1995; 33:237–56.

39. Infante-Malachias ME, Yotoko KSC, Lima De Azeredo Espin AM. Random amplified polymorphic DNA of screwworm fly populations (Diptera: Calliphoridae) from Southeastern Brazil and Northern Argentina. Genome 1999; 42:772–9.

40. Lyra ML, Fresia P, Gama S, Cristina J, Klaczko LB, de Azeredo-Espin AML. Analysis of Mitochondrial DNA Variability and Genetic Structure in Populations of New World Screwworm Flies (Diptera: Calliphoridae) from Uruguay. J Med Entomol 2005; 42:589–95.

41. Torres TT, Lyra ML, Fresia P, Azeredo-Espin AML. Assessing Genetic Variation in New World Screwworm *Cochliomyia hominivorax* Populations from Uruguay [Internet]. In: Vreysen MJB, Robinson AS, Hendrichs J, editors. Area-Wide Control of Insect Pests. Dordrecht: Springer Netherlands; 2007 [cited 2021 Oct 27]. page 183–91. Available from: http://link.springer.com/10.1007/978-1-4020-6059-5_16

42. Lyra ML, Klaczko LB, Azeredo-Espin AML. Complex patterns of genetic variability in populations of the New World screwworm fly revealed by mitochondrial DNA markers. Medical Vet Entomology 2009; 23:32–42.

43. Torres TT, Azeredo-Espin AML. Population genetics of New World screwworm from the Caribbean: insights from microsatellite data. Medical Vet Entomology 2009;23:23–31.

44. Fresia P, Lyra ML, Coronado A, De Azeredo-Espin AML. Genetic Structure and Demographic History of New World Screwworm Across Its Current Geographic Range. jnl. med. entom. 2011; 48:280–90.

45. Tietjen M, Pérez De León AA, Sagel A, Skoda SR, Phillips PL, Mitchell RD, et al. Geographic Population Genetic Structure of the New World Screwworm, *Cochliomyia hominivorax* (Diptera: Calliphoridae), Using SNPs. Journal of Medical Entomology 2022; 59:874–82.

46. Torres TT, Brondani RPV, Garcia JE, Azeredo-Espin AML. Isolation and characterization of microsatellite markers in the new world screwworm *Cochliomyia hominivorax* (Diptera: Calliphoridae). Molecular Ecology Notes 2004; 4:182–4.

47. Torres TT, De Azeredo-Espin AML. Development of new polymorphic microsatellite markers for the New World screwworm *Cochliomyia hominivorax* (Diptera: Calliphoridae). Molecular Ecology Notes 2005; 5:815–7.

48. Nei M. Estimation Of Average Heterozygosity And Genetic Distance From A Small Number Of Individuals. Genetics 1978; 89:583–90.

49. Goudet J. Hierfstat, a package for R to compute and test hierarchical *F*-statistics. Molecular Ecology Notes 2005; 5:184–6.

50. Raymond M, Rousset F. An Exact Test For Population Differentiation. Evolution 1995; 49:1280–3.

51. Rousset F. GENEPOP ‘007: a complete re-implementation of the GENEPOP software for Windows and Linux. Molecular Ecology Resources 2008; 8:103–6.

52. Piry S, Luikart G, Cornuet JM. Computer note. BOTTLENECK: a computer program for detecting recent reductions in the effective size using allele frequency data. Journal of Heredity 1999; 90:502–3.

53. Weir BS, Cockerham CC. Estimating F-Statistics for the Analysis of Population Structure. 1984.

54. Goudet J. FSTAT (Version 1.2): A Computer Program to Calculate F-Statistics. Journal of Heredity 1995; 86:485–6.

55. Rice WR. Analyzing Tables of Statistical Tests. Evolution 1989; 43:223.

56. Rousset F. Genetic Differentiation and Estimation of Gene Flow from *F* -Statistics Under Isolation by Distance. Genetics 1997; 145:1219–28.

57. R Core Team. R: A Language and Environment for Statistical Computing. [Internet]. 2023; Available from: https://www.R-project.org/

58. Waples RS. Testing for Hardy–Weinberg Proportions: Have We Lost the Plot? Journal of Heredity 2015; 106:1–19.

59. Wahlund S. Zusammensetzung von populationen und korrelationserscheinungen vom standpunkt der vererbungslehre aus betrachtet. Hereditas 2010; 11:65–106.

60. Zhivotovsky LA. Relationships Between Wright’s FST and FIS Statistics in a Context of Wahlund Effect. Journal of Heredity 2015; 106:306–9.

61. Dharmarajan G, Beasley JC, Rhodes OE. Heterozygote deficiencies in parasite populations: an evaluation of interrelated hypotheses in the raccoon tick, Ixodes texanus. Heredity 2011; 106:253–60.

62. Garnier-Géré P, Chikhi L. Population Subdivision, Hardy–Weinberg Equilibrium and the Wahlund Effect [Internet]. In: Encyclopedia of Life Sciences. Wiley; 2013 [cited 2024 Apr 29]. Available from: https://onlinelibrary.wiley.com/doi/10.1002/9780470015902.a0005446.pub3

63. De Meeûs T. Revisiting FIS, FST, Wahlund Effects, and Null Alleles. Journal of Heredity 2018; 109:446–56.

64. Gaggiotti OE, Bekkevold D, Jørgensen HBH, Foll M, Carvalho GR, Andre C, et al. Disentangling the effects of evolutionary, demographic, and environmental factors influencing genetic structure of natural populations: atlantic herring as a case study. Evolution 2009; 63:2939–51.

65. Baqir HA, Ab Majid AH. Population genetic structure of tropical bed bug (Hemiptera: Cimicidae) populations and their breeding pattern in Iraq. Journal of Insect Science 2024; 24:9.

66. Cockerham CC, Weir BS. Estimation of Gene Flow from F-Statistics. Evolution 1993; 47:855.

67. Ma L, Ji YJ, Zhang DX. Statistical measures of genetic differentiation of populations: Rationales, history and current states. Current Zoology 2015; 61:886–97.

68. Donnelly MJ, Licht MC, Lehmann T. Evidence for Recent Population Expansion in the Evolutionary History of the Malaria Vectors Anopheles arabiensis and Anopheles gambiae. Molecular Biology and Evolution 2001; 18:1353–64.

69. Fountain T, Duvaux L, Horsburgh G, Reinhardt K, Butlin RK. Human-facilitated metapopulation dynamics in an emerging pest species, *C imex lectularius*. Molecular Ecology 2014; 23:1071–84.

70. Walt HK, King JG, Sheele JM, Meyer F, Pietri JE, Hoffmann FG. Do bed bugs transmit human viruses, or do humans spread bed bugs and their viruses? A worldwide survey of the bed bug RNA virosphere. Virus Research 2024; 343:199349.

71. Lehmann T, Hawley WA, Kamau L, Fontenille D, Simard F, Collins FH. Genetic differentiation of *Anopheles gambiae* populations from East and West Africa: comparison of microsatellite and allozyme loci. Heredity 1996; 77:192–200.

72. Besansky NJ, Lehmann T, Fahey GT, Fontenille D, Braack LEO, Hawley WA, et al. Patterns of Mitochondrial Variation Within and Between African Malaria Vectors, *Anopheles gambiae* and *An. arabiensis*, Suggest Extensive Gene Flow. Genetics 1997; 147:1817–28.

73. Bonizzoni M, Zheng L, Guglielmino CR, Haymer DS, Gasperi G, Gomulski LM, et al. Microsatellite analysis of medfly bioinfestations in California. Molecular Ecology 2001; 10:2515–24.

74. Baliraine FN, Bonizzoni M, Guglielmino CR, Osir EO, Lux SA, Mulaa FJ, et al. Population genetics of the potentially invasive African fruit fly species, Ceratitis rosa and Ceratitis fasciventris (Diptera: Tephritidae). Mol Ecol 2004; 13:683–95.

75. Agis M, Schlotterer C. Microsatellite variation in natural Drosophila melanogaster populations from New South Wales (Australia) and Tasmania. Mol Ecol 2001; 10:1197–205.

76. Roslin T. Spatial population structure in a patchily distributed beetle: POPULATION STRUCTURE IN *APHODIUS FOSSOR*. Molecular Ecology 2001; 10:823–37.

77. Felius M, Beerling ML, Buchanan D, Theunissen B, Koolmees P, Lenstra J. On the History of Cattle Genetic Resources. Diversity 2014; 6:705–50.

78. Dear JP. A revision of the New World Chrysomyini (Diptera: Calliphoridae). Rev. Bras. Zool. 1985; 3:109–69.

79. Hendry AP, Day T. Population structure attributable to reproductive time: isolation by time and adaptation by time. Molecular Ecology 2005; 14:901–16.

80. Duforet-Frebourg N, Slatkin M. Isolation-by-distance-and-time in a stepping-stone model. Theoretical Population Biology 2016; 108:24–35.

81. Owings CG, Banerjee A, Picard CJ. Temporal population genetic structure of *Phormia regina* (Diptera: Calliphoridae). Journal of Medical Entomology 2023; tjad115.

82. Peniston JH, Backus GA, Baskett ML, Fletcher RJ, Holt RD. Ecological and evolutionary consequences of temporal variation in dispersal. Ecography 2024; 2024:e06699.

83. Dobson A. Metalife! Science 2003; 301:1488–90.

84. Hanski IA, Gaggiotti OE. Ecology, Genetics and Evolution of Metapopulations. Saint Louis: Elsevier Science; 2014.

85. Phillips PL, Welch JB, Kramer M. Seasonal and Spatial Distributions of Adult Screwworms (Diptera: Calliphoridae) in the Panama Canal Area, Republic of Panama. J Med Entomol 2004; 41:121–9.

86. Israely N, Ziv Y, Galun R. Metapopulation Spatial–Temporal Distribution Patterns of Mediterranean Fruit Fly (Diptera: Tephritidae) in a Patchy Environment. an 2005; 98:302–8.

87. Ke F, Li J, Vasseur L, You M, You S. Temporal sampling and network analysis reveal rapid population turnover and dynamic migration pattern in overwintering regions of a cosmopolitan pest. Front. Genet. 2022; 13:986724.

88. Taylor CE, Manoukis NC. Effective population size in relation to genetic modification of Anopheles gambiae sensu stricto. In: Takken W, Scott W, editors. Ecological aspects for application of genetically madified mosquitoes. Kluwer Academic Publishers, Dordrecht.; 2002. page 133–46.

89. Ibrahim KM. Plague dynamics and population genetics of the desert locust: can turnover during recession maintain population genetic structure? Molecular Ecology 2001; 10:581–91.

90. Mendonça BAA, De Sousa ACB, De Souza AP, Scarpassa VM. Temporal genetic structure of major dengue vector Aedes aegypti from Manaus, Amazonas, Brazil. Acta Tropica 2014; 134:80–8.

